# Whole genome screening defines a key role of autophagy in resistance of bovine cells to BVDV infection

**DOI:** 10.64898/2026.03.24.712903

**Authors:** Christiane Riedel, Hann-Wei Chen, Till Rümenapf, Florian Grebien, Maren van Son, Thomas Nelson Harvey, Matthew Peter Kent, Victor Boyartchuk

**Affiliations:** CIRI (Centre International de Recherche en Infectiologie), Univ Lyon, Inserm, U1111, Université Claude Bernard Lyon 1, CNRS, UMR5308, ENS de Lyon, 69007 Lyon, France; Institute of Virology, University of Veterinary Medicine, Vienna, Austria; Institute of Biochemistry, University of Veterinary Medicine, Vienna, Austria; Norsvin SA, Hamar, Norway; Center for Integrative Genetics (CIGENE), Faculty of Biosciences (BIOVIT), Norwegian University of Life Sciences, Aas, NO-1432, Norway

## Abstract

Bovine viral diarrhea virus (BVDV, genus *Pestivirus*, family *Flaviviridae*) is a notifiable pathogen of cattle which significantly impacts animal health, welfare, and the economy. Several cellular factors important for BVDV infection, such as Jiv, CD46 and ADAM17, have already been identified providing new targets development of effective defense strategies. However, our knowledge about BVDV host factor requirements remains limited, as no genome-wide studies of BVDV host resistance factors were performed to date, in part due to lack of accessible whole genome libraries. To close this gap, we have designed a novel bovine whole genome knockout library and successfully used it to identify a set of BVDV host resistance factors. The validity of our approach is highlighted by the strong selection of cells with inactivated ADAM17 and TMEM41B, which have both been described to be of pivotal importance for BVDV infection. In addition, guides targeting VMP1, recently identified as an important factor for flavivirus infection, were also significantly enriched in our screen. Furthermore, we found differential selection of several proteins essential for triggering autophagy, providing additional strong evidence of this process underlying key cellular functions involved in resistance to BVDV.

## Introduction

In the last decade CRISPR-Cas9 based whole genome screening has greatly accelerated the discovery of genes controlling a wide range of phenotypes, including susceptibility to viral infections (1). This approach has been extensively used to study the susceptibility of human and murine cells to different pathogens, enabled in large by availability of validated whole genome libraries (2–4). Similar studies in cellular models of economically relevant production animal pathogens lag behind due to the lack of such tools. For cattle, only recently reports of development and validation of whole genome libraries have begun to emerge (5, 6)

Bovine viral diarrhea virus (BVDV) is a small, enveloped, single-stranded, positive-sense RNA virus within the genus *Pestivirus* (7) (family *Flaviviridae*). It is the causative agent of bovine viral diarrhea, which is listed as a reportable disease by the World Organization for Animal Health (8). Despite mandatory control and eradication programs implemented by most European countries, it continues to pose a notable threat to animal welfare and results in significant negative economic impact (9, 10). BVDV often persists in animals that were infected *in utero* as a result of immunotolerance (11). Such carriers pose a significant epidemiological threat, as they remain asymptomatic while shedding large amounts of virus (12).

The economic and animal welfare impacts of BVDV drive sustained efforts towards the discovery of targetable host factors that could be used to modulate susceptibility to this infection. To date, several genes have been characterized as essential for BVDV entry and replication. The cell surface metalloprotease ADAM17 has been shown to serve as a key viral entry factor and cells lacking ADAM17 are completely resistant to BVDV infection (13). Another cell surface protein, CD46 has been identified as a cellular receptor for BVDV (14). It is currently being actively pursued a key candidate for creating BVDV resistant cattle (15).

The BVDV genome has a length of approximately 12.5kb and its single open reading frame encodes one polyprotein, which is co- and post-translationally processed by cellular and viral proteases into at least 12 mature proteins. Pestiviruses are masters of evading detection by the host’s innate immune response, for example by inhibiting the central transcription factor IRF3 and preventing the detection of their RNA genomes by toll like receptors. BVDV is classified into two distinct biotypes based on its effect on infected cells. The predominant non cytopathic biotype depends on the cellular chaperone DNAJC14 to promote viral NS2 autoprotease activity, and its absence renders cells non-permissive for BVDV infection. On the contrary, the cytopathic biotype, which is usually isolated from cases of mucosal disease in immunotolerant cattle, has acquired modifications of the genome rendering NS2-3 cleavage DNAJC14 independent (16). Another viral factor capable of modulating host cellular processes is the non-structural protein 4B (NS4B). It has been shown to promote autophagy in bovine MDBK cells (17), suggesting that the virus actively uses this compartment for its persistence.

The intricate interplay and dependence of BVDV on host factors is also highlighted by recent genome wide KO screens aiming to identify essential host factors for infection with Zika and Yellow Fever flaviviruses (3). This study identified TMEM41B and VMP1, scramblases (18, 19) as important for the regulation of ER membrane dynamics and autophagy (20). The subsequent testing of 14 families of flaviviruses, including BVDV defined TMEM41B as pan-flavivirus host factor (3).

Autophagy is a fundamental evolutionary conserved process that was originally discovered as a recycling mechanism for breakdown, removal, and reuse of unneeded biomolecules (21). Subsequently, it has been recognized that it is essential for cell survival under a wide range of stress-inducing conditions, including infections with viral pathogens. Generally, cells use autophagy to restrict viral replication, which provides selective pressure for a number of viruses (e.g. *Herpesviruses*, HIV-1) to evolve mechanisms to inhibit this process (22). As an alternative host avoidance mechanism, a subset of viruses, including *Flaviviruses*, use autophagic compartments as a protected environment for replication and maturation (23).

Considering challenges posed by the complexity of BVDV infections and lack of accessible whole genome manipulation tools we set out to develop a novel bovine whole genome knock-out library and use it to identify host factors controlling resistance to the virus.

## Results

### Novel whole genome CRISPR knockout library

To enable systematic discovery of genes that play key roles in resistance of bovine cells to BVDV we have designed and constructed a novel whole genome CRISPR Cas9 knockout library using an in-house assembled pipeline for selection, scoring, and assembly of sgRNA guides. Our goal was to target each coding and lncRNA gene with five individual guides. Due to specific constraints imposed on the type of sequences that can serve as guides for the CRISPR-Cas9 nuclease (location within transcribed sequences, adjacent NGG etc.) the total number of coding genes targeted by five guides was 21,071, or 96% of the genes annotated in the v104 Ensembl ARS-UCD1.2 *Bos taurus* genome assembly (Sup Fig 1A). Out of 1488 annotated lncRNAs our library targets 1459 with five guides (Sup Fig 1B), with only eight lncRNA lacking any targeting guides. Following the library synthesis, guides were cloned into the custom pLV2.8 lentiviral vector derived by replacing the puromycin resistance gene of pKLV2-U6gRNA5(BbsI)-PGKpuro2ABFP-W (24) with the sequence encoding the blasticidin resistance gene. The overall representation of guides in the cloned library was measured by NGS and found to be largely uniform, with 70% of guide frequencies residing within 1 SD (9.4×10^−12^) from the expected value of 8.77×10^−6^ percent (Sup Fig 1C). Only three of the designed guides were not found in the final cloned library. Based on this analysis of guide representation in the cloned library, we conclude that our library has good coverage of currently annotated genes.

**Fig 1.**
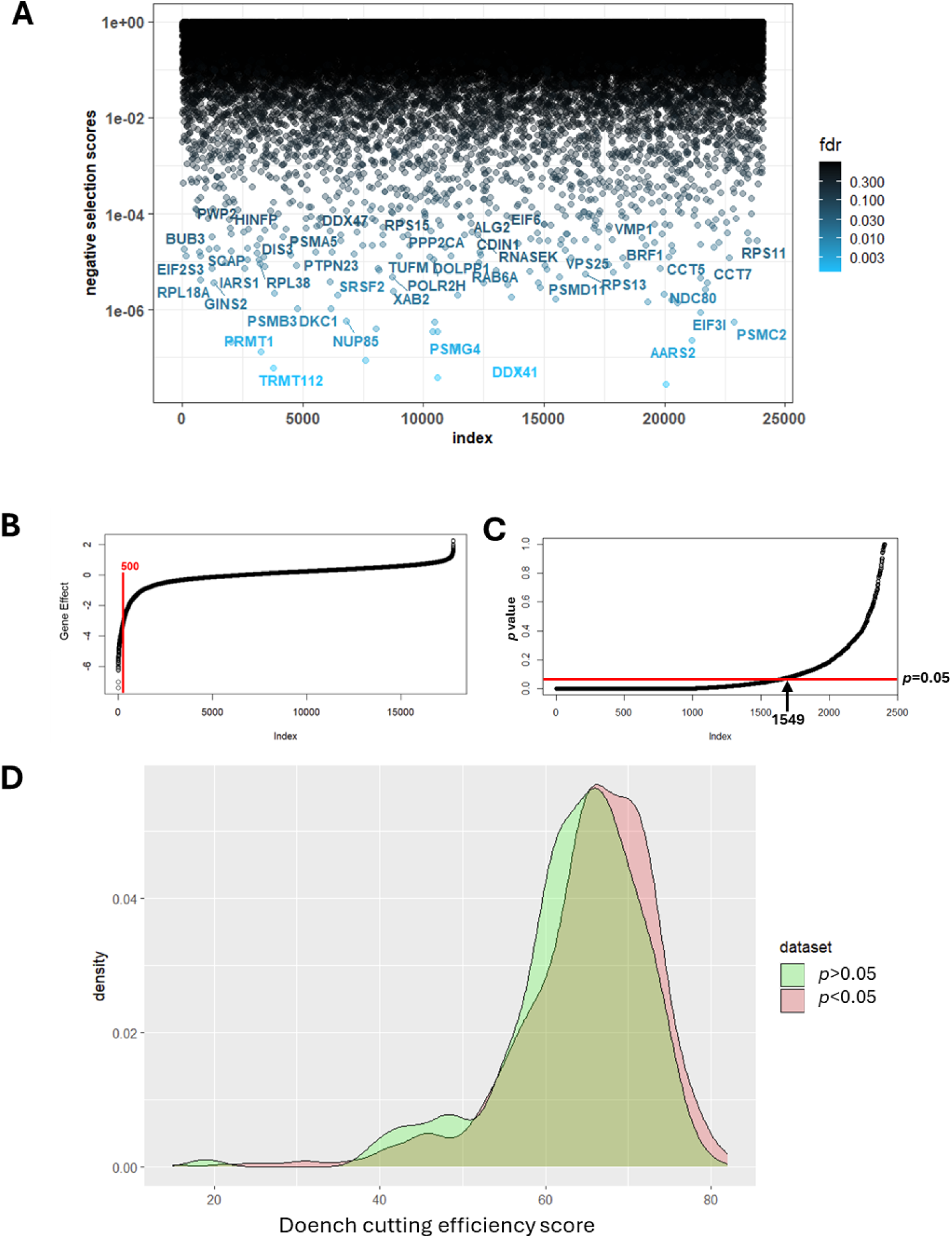
Evaluation of the novel bovine whole genome knockout library. (**A**) 352 essential genes were significanly depleted (FDR<0.1) in MDBK cells grown for 22 days after the library transduction. (**B**) Distibution of combined human kidney cell line gene effects used to define a set of 500 essential genes. (**C**) *p*-values of observed decrease in representation of individual guides (five per gene) targeting bovine homologues of the selected 500 human essential genes. (**D**) Distributions of Doench cutting efficiency scores of guides that were significanly (p<0.05) depleted in the Day 22 population as well as guide whose representation did not significntly change (p> 0.05).

### Establishment of Cas9 expressing MDBK cells

To facilitate genome wide screening experiments, we created an MDBK cell line inducibly expressing Cas9 and GFP under the control of the Tet responsive element. These cells were generated by co-transfection of existing MDBK Tet-On cells with a plasmid encoding the Cas9-P2A-GFP cassette and an additional plasmid carrying a puromycin resistance gene to allow for subsequent antibiotic selection. The resulting clonally selected, puromycin resistant cell population readily showed expression of GFP after stimulation with Doxycycline. The functionality of Cas9 was assessed by quantifying the decrease of GFP fluorescence after expression of an sgRNA targeting GFP. Already 3 days after sgRNA expression, more than 70% of the cells showed diminished GFP levels, whilst cells transduced with an empty sgRNA expression vector remained nearly 100% GFP positive as determined by flow cytometry (Sup Fig 2A). Further confirmation of Cas9 activity was obtained by targeting the essential gene *myc* using an sgRNA delivered by a vector also expressing mCherry. A decline in mCherry expressing cells could be observed starting 10 days after transduction, whilst the amount of mCherry expressing cells treated with the control plasmid lacking sgRNA remained at the same level (Sup Fig 2A). With the functionality of Cas9 established, the infectability of the MDBK_Cas9_ cells by BVDV was compared with the parental MDBK cells. When using defined virus dilutions, no difference in the amount of detected infectious virus was identified between the two cell lines, indicating that virus entry and replication are comparably efficient (Sup Fig 2B). Therefore, the newly generated MDBK_Cas9_ cells were considered an appropriate model to use in the screen aimed at the identification of BVDV host factor requirements.

**Fig 2.**
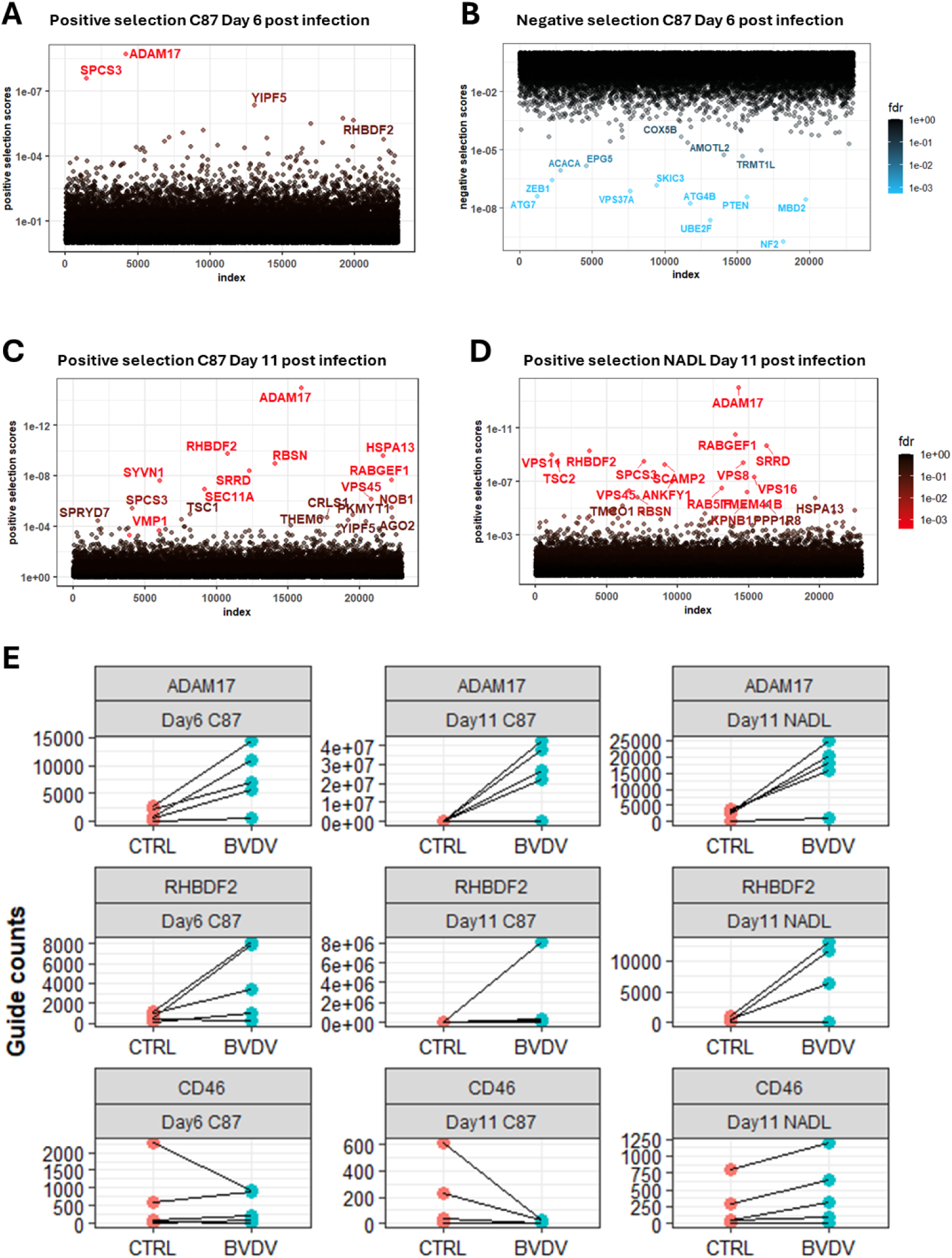
Genome-wide screen BVDV resistance screen identifies host factors required for resistance. (**A**) Bubble plot of positively and (**B**) negatively selects gene knockouts after 6-day exposure to BVDV C87_mCherry-E2_. By Day 11 post infection there were almost no identifiable negatively selected mutans, while the number of positively selected knockouts substantially increased for both the C87_mCherry-E2_ (**C**) and NADL (**D**) infected populations. Normalized counts of individual guides targeting BVDV susceptibility factors ADAM 17 and RHBDF2 increased following viral infections, while no significant changes were detected for guides targeting CD46 (**E**).

### Performance of the novel bovine library

Following transduction of our knockout library and selection with blasticidin for MDBK_Cas9_ cells containing the sgRNA expressing vector, we quantified relative guide abundance in the resulting cell population on day 22 after transduction (Fig. 1A). When compared to the individual guide frequency in the original cloned library, we observed a significant (FDR<0.1) decrease in abundance of guides targeting 352 genes. Such depletion is expected and likely results from death of cells expressing guides targeting genes essential for viability (Sup. Fig 1D). While overall such a depletion reduces the gene space that can be interrogated in a knockout screen, it can be useful as a quality control measure for the performance of our library. This can be done by comparison of observed depleted genes to previously defined essential gene sets. Such comparisons are straightforward to obtain since extensive cell line specific gene essentiality information is publicly available, especially for human cells or cells from model organisms. Human cell line specific screening datasets are available for download on the DepMap cancer dependency map project portal (https://depmap.org/portal/). To obtain an overview of kidney cell specific essential genes, we averaged z-ranks of essentiality scores from three human immortalized embryonic kidney cell lines: HA1E, HK2 and HEKTE (25). We have arbitrarily chosen to use 500 genes with the lowest ranks to create a benchmark set of genes that control viability of immortalized human kidney cells (Fig 1B). We then used this set to check how many of our library guides targeting bovine homologues were underrepresented. Out of 500 human genes there were 491 genes that had annotated bovine homologues, and our library contained 2404 out of possible 2455 guides targeting these homologues. In the day 22 dataset, the abundance of 1549 (64% of total) guides targeting 491 bovine homologues was significantly (p<0.05) lower than in the original library (Fig. 1C). To determine if there are any quantifiable differences between guides that were depleted and those that were not, we plotted distributions of Doench cutting efficiency score (26) that was initially computed for each guide (Materials & Methods) and used them as a factor for inclusion in the library (Fig 1D). We found that while distributions were largely overlapping, there was a notable and statistically significant shift towards the lower Doench scores for guides that were not significantly depleted. This observation indirectly confirms the utility of the cutting efficiency scores even for organisms that were not included in the original model training sets.

### Mutations in ADAM17 and RHBDF2 are enriched in populations surviving BVDV challenge

To test the importance of host factors in survival-based screening, we chose to work with two cytopathic BVDV strains, the type 1a strain NADL and an in-house generated, lower virulence C87_mCherry-E2_ strain, that is tagged with mCherry (27). We chose cytopathic BVDV strains in our experimental setup to facilitate the selection of infected cells. Hence, this choice renders our screen unable to identify cellular factors involved in DNAJC14 dependent NS2-3 cleavage but should otherwise serve as an adequate model for host factor requirements of non-cytopathic BVDV isolates. To initiate the screen, the transduced cell populations under blasticidin selection were infected with either NADL or C87_mCherry-E2_ with an MOI of 0.1, respectively. Genomic DNA was recovered from surviving cells after six (for C87_mCherry-E2_) and eleven (for C87_mCherry-E2_ and NADL) days, as well as from uninfected control populations cultured in parallel for the same period of time and used to create PCR amplicon libraries of guides remaining in each population. After six days of infection with C87_mCherry-E2_, short read Illumina sequencing and bioinformatic comparison of control and treated cell guide frequencies identified significant positive (Fig 2A) and negative (Fig 2B) enrichments in guide abundance. As expected, surviving treated cells showed a significant positive enrichment for guides targeting the essential BVDV entry factor ADAM17, indicating that mutations in this gene (leading to its inactivation) offer protection from C87_mCherry-E2_ induced cell death (13). Deletions in the gene encoding the endoplasmic reticulum resident regulator of ADAM17 biogenesis, *RHBDF2* aka iRhom2, were also overrepresented in day six population. Even though the direct involvement of RHBDF2 in the control of BVDV infections has not been reported, this gene acts as a binding factor and upstream regulator of ADAM17 (28) providing direct explanation of its role in BVDV infection.

Eleven days post-infection the number of positively selected (FDR<0.1) mutants in C87_mCherry-E2_ (Fig 2 C) challenged populations had increased compared to day six (8 at D6 vs 22 at D11), with even more being detected in NADL at eleven days (n=58; Fig 2D). For both viral strains, *ADAM17* and *RHBDF2* mutants remained among the most overrepresented mutants. Interestingly, an overlap of only eight genes was observed between the 19 highly significantly enriched (FDR<0.05) genes observed for C87_mCherry-E2_ infection and the 35 observed for NADL (Sup Table 2)

Surprisingly, in MDBK knockout cell populations challenged with either C87_mCherry-E2_ or NADL, no positive selection for *CD46* mutants was detected. CD46 is a receptor that mediates BVDV entry, and it has recently been shown that mutations in this gene reduce the susceptibility of bovine cells to infection by 90-99% (29). We have confirmed the presence of individual guides at each timepoint but, in contrast to *ADAM17* or *RHBDF2* targeting guides (Fig 2E), their abundance did not significantly change in infected populations. In the ranked list of genes with positive enrichment of guides, CD46 occupies position 515 for infection with NADL, and even lower ranks (2,323 for D6 and 6,917 for D11) for C87_mCherry-E2_. This difference in ranks is likely due to the approximately 10-fold higher dependence of NADL on CD46 (30). Furthermore, it also provides an idea about the sensitivity of the screen to identify factors with lower impact on BVDV infection.

### Autophagy is a key process controlling the outcome of BVDV infection

To identify common cellular functions that were impacted by gene deletions that were differentially selected following infection with BVDV, we performed gene set enrichment analysis. Gene Ontology term analysis of the top 40 genes that were differentially selected by day six post infection with C87_mCherry-E2_ identified enrichment for various aspects of peroxisomal and vacuolar trafficking (Fig 3A). Curated Reactome (31) pathway gene set analysis provided further support for the involvement of peroxisomes (Fig 3B). Interestingly, the Reactome analysis also pointed towards autophagy as a process controlling BVDV infection. This was further supported by KEGG pathway analysis (32) of the same set of differentially selected genes that revealed autophagy as the only pathway that was significantly enriched (Fig 3C and Sup Fig 3). Based on these findings we chose to focus our screen result validation tests on genes involved in cellular autophagy.

**Fig 3.**
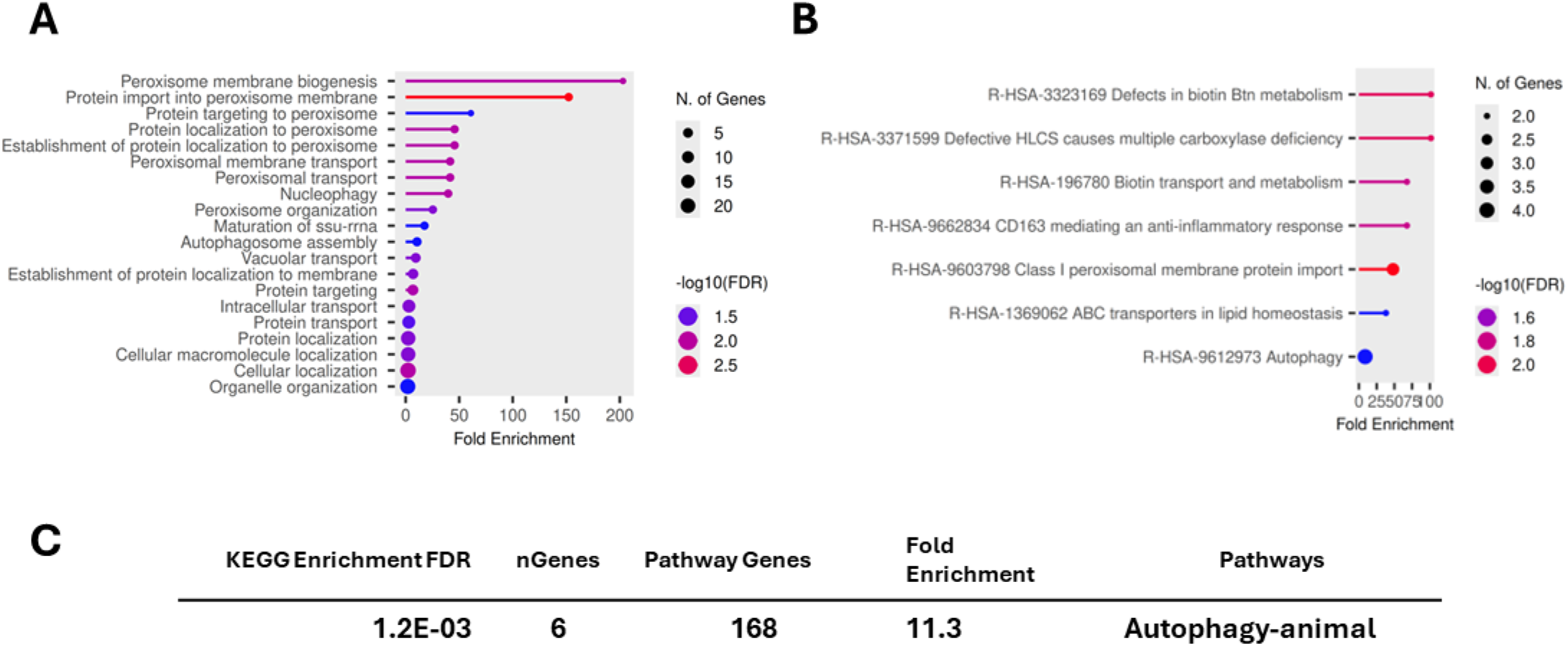
Gene set and pathway enrichment analysis using top 44 differentially selected genes. Significantly enriched Gene Ontology Molecular Function terms (**A**) point to the role of peroxisomes in resistance to infection. Biotin biogenesis and inflammation Reactome pathway gene sets are highly enriched by the selected genes (**B**). KEGG pathway analysis results (**C**) reveal autophagy as the only significantly enriched pathway, consistent with the results of GO and Reactome analysis

### Survival of BVDV infected cells lacking genes controlling autophagy

To serve as a technical validation of the screen and confirm their role in BVDV pathogenesis, we chose a set of six genes for follow up in single gene knockout BVDV challenge experiments (Table 1). Five of the six selected genes have documented involvement in autophagy, of which four were negatively selected in the screen, while *VMP1* was under positive selection. We also included *PHGDH* based on recent reports documenting its involvement in innate immune control of classical swine fever infection (33). To generate verification knockouts, we used two different guides for each gene; one selected from among the five used in the screen (L_ guide, Sup Table 1) and one new design that was not included in the original library (C_ guide). To exclude the possibility that our phenotypes were artifacts only observable in our MDBK_Cas9_ cells, these control experiments were performed in the parental MDBK cells transduced with a lentiviral vector expressing gRNA from the U6 promoter and Cas9 from the EF-1a core promoter (LentiCRISPRv2-Opti). To avoid issues related to multiple lentiviral integrations in the host genome, our transductions were performed at low MOI (<0.5). This creates a situation where cells receive a variable, but generally low, number of integrations. However, in some cases puromycin, expressed from a separate promoter, can be produced at sufficient levels to maintain selection, while either Cas9 or guide expression levels being too low, resulting in incomplete editing of the targeted locus. Instead of performing single cell cloning to create 100% edited populations, we chose to capitalize on this mixed population of wild type and sgRNA expressing cells. We proceeded to quantify the proportion of knock out cells in challenged and control cells using bioinformatic deconvolution of Sanger sequencing traces of the edited loci using the *ice* bioinformatic pipeline (34). In two independent experiments we have challenged mixed cell populations with either C87_mCherry-E2_ or NADL for either five or seven days. Genomic DNA was then isolated from challenged individual gene knockout populations, and the control population grown in parallel. Following PCR amplification of the targeted locus, it was sequenced using an internal primer and the resulting sequencing trace deconvoluted. For almost all tested genes we were able to obtain at least two independent results confirming their involvement in the control of BVDV infection (Table 2). For example, in two independent experiments using two different guides, cells exposed to C87_mCherry-E2_ had an up to 3-fold decrease in the proportion of cells with guides targeting the *ATG4B* gene (Fig 4A). The proportion of *ATG4A* gene modifications in cells infected with NADL also decreased, but to a smaller extent. For both genes in our set that were targeted by positively selected guides in the screen (*VMP1* and *PHGDH*), we saw a dramatic increase in the proportion of edited cells following challenge with BVDV (Table 2). As shown in Figure 4B for *VMP1* targeting guide C_VMP1, there were no knockout cells detected by the deconvolution software in the unchallenged population. This is likely due to *VMP1* being an essential gene causing mutant cells to be rapidly outcompeted. However, after challenge with BVDV, there was a significant increase in the proportion of mutant cells for both *VMP1* targeting guides. Interestingly NADL was able to drive the selection of *VMP1* mutants to a much higher extent than C87_mCherry-E2_.

**Fig 4.**
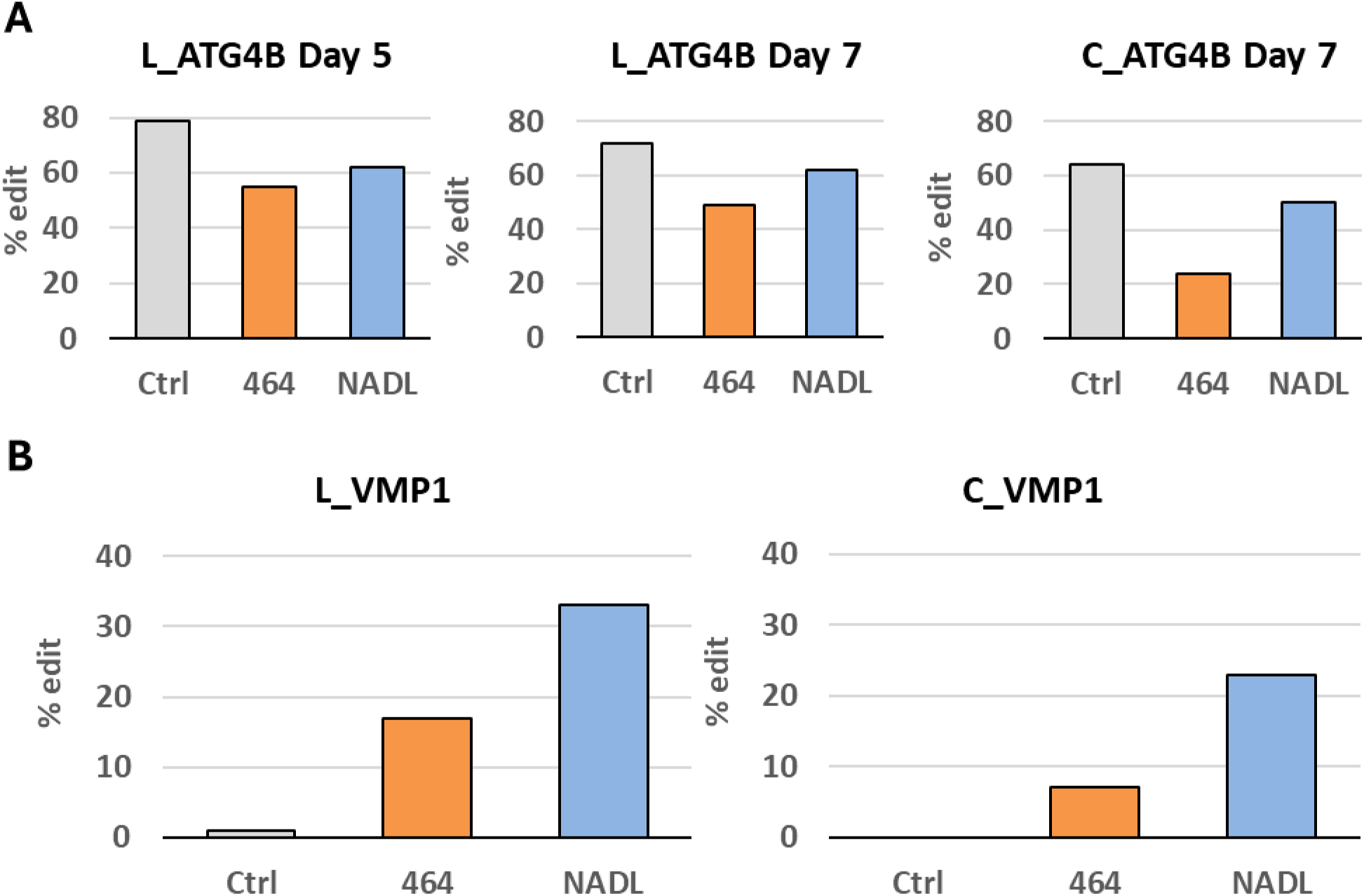
Percentage of edited cells in control and BVDV infected populations. BVDV infection depletes ATG4B mutants (**A**) created using two different guides (L_ sgRNA that was included in the original library and C_ sgRNA that was distinct from the five library guides) in two independent experiments. BVDV infection selects for VMP1 loss of function resulting in a dramatic increase in the proportion of VMP1 mutants (**B**) in challenged populations.

**Table 1.** Genes used in single gene knockout validation experiments.

| Gene | Ensembl id | LFC | *p* value | FDR | score |
| --- | --- | --- | --- | --- | --- |
| ATG4B | ENSBTAG00000015401 | -7.0206 | 2.07E-07 | 0.00055 | 1.76E-08 |
| ATG7 | ENSBTAG00000035827 | -4.1166 | 2.07E-07 | 0.00055 | 3.91E-08 |
| ATG3 | ENSBTAG00000009084 | -4.557 | 7.09E-05 | 0.099884 | 2.30E-05 |
| ATG12 | ENSBTAG00000017525 | -1.5298 | 1.27E-03 | 0.486709 | 3.80E-04 |
| VMP1 | ENSBTAG00000011623 | 5.2139 | 2.75E-05 | 0.086015 | 6.22E-06 |
| PHGDH | ENSBTAG00000039719 | 5.3073 | 1.96E-05 | 0.086015 | 1.83E-06 |

**Table 2.**
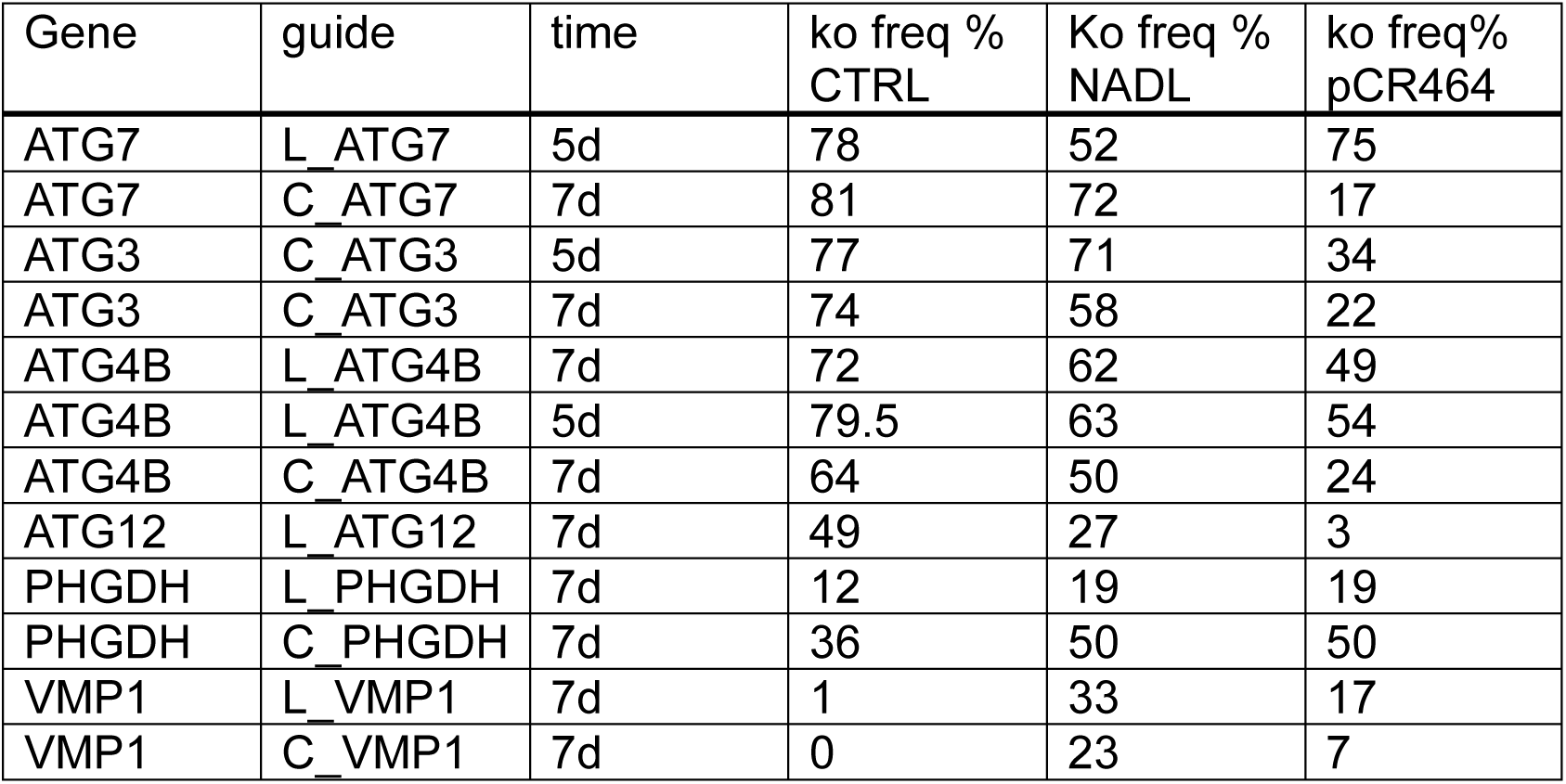
Summary of independent tests mutation frequency in BVDV challenged single gene knockouts.

Overall, the population-based analysis of individual knock outs confirmed the main finding of our screen that autophagy plays a key role in the control of BVDV infection of bovine cells.

## Discussion

Pestiviruses remain a significant threat to the domestic animal population and cause significant economic losses and animal welfare problems. Hence, a strong interest remains to identify host genes that can lower the susceptibility of farm animals to pestivirus infection, through either breeding efforts or genetic modification. In this context, detailed knowledge of host factors important for pestivirus infection is pivotal.

Whole genome mutant screens provide a powerful platform for unbiased discovery of key genetic elements controlling an observed phenotype. For a long period of time such screens were limited to random mutagenesis of primarily haploid organisms. However, the development of genome-scale CRISPR-Cas9 KO screening (35, 36) now allows to conduct such screens in any organism with a sequenced genome. The majority of whole genome screens to date have been conducted in human or rodent model cells. This is in large part due to early development, validation and public availability of human and mouse whole genome libraries and Cas9 expressing cell lines. Similar studies in non-model organisms, such as economically important production animals, are rare due to a lack of such resources. Only recently, two screens developed in bovine cells were published. These libraries consist of 89,566 (37) or 94,000 guides (5) targeting 21,241 or 21,165 genes respectively.

To increase the gene space sampled and include guides for some of the regulatory sequences, we chose to design and construct our own library, which targets 21,484 annotated protein coding genes and 1,480 long non-coding regions, using a total of 113,024 specific, and 1,000 control, guides. A combination of publicly available bioinformatic tools were used to evaluate each potential guide for its on- and off-target activity. To estimate the performance of the chosen guides, we analyzed the depletion of guides targeting putative essential genes, using information of gene essentiality available for human kidney cell lines. These tests indicate that the bioinformatic pipeline used for library design reliably assessed the cutting efficiency also in non-model organisms.

In most whole genome screens sgRNA guides are either introduced individually into cells expressing Cas9 nuclease or delivered together with a gene coding for Cas9 cloned in the same construct as the guide. Cas9 expressing cells provide the benefit of uniform expression of the editing nuclease thus removing one of factors affecting efficiency of the edits. In our study we chose to generate a MDBK cell line inducibly expressing Cas9 and GFP under the control of the *tet* responsive element, to limit negative effects of continuous Cas9 expression on the cells (38).

The performance of the newly designed library and the reliability of its results were further strengthened by the identification of several factors with an already known key role in BVDV infection, namely the cell surface sheddase ADAM17, also known as TACE, and the ER membrane protein TMEM41B (3, 13). In addition to these two genes, we identified two additional genes with a likely impact on BVDV infection, namely the essential *ADAM17* trafficking factor, iRhom2, as well as the Vacuole Membrane protein *VMP1*. VMP1 has been previously identified as a host factor used for viral replication in the pan-flavivirus screen, but it was not tested for its ability to mediate BVDV infection. Interestingly, our screens did not provide any evidence of the selective advantage of cells with mutated *CD46*. CD46 is a long known, non-essential BVDV receptor, deletion of which results in a 5- to100-fold decrease of cellular susceptibility to infection with BVDV (30, 39, 40). Whilst this reduction of cellular susceptibility is of biological relevance, for example by preventing BVDV related disease in a genetically modified calf (41), the effect of *CD46* deletion is relatively low when compared to the effect of the ADAM17 KO, which results in complete cellular resistance to BVDV infection (13). This difference in strength of infection inhibition could explain the lack of detection of a selective advantage of *CD46* mutants in our survival-based screen. To further assess this possibility, sorting infected cells by flow cytometry according to the levels of virus replication could provide an insight if such an approach is advantageous over a survival-based screen in identifying candidates with lower biological impact.

Autophagy is an essential cellular process that handles a wide range of tasks, from recycling unwanted biomolecules to defending against invading pathogens. Flaviviruses evolved to take advantage of this process by using its compartments for replication and immune evasion (42) with several studies highlighting the importance of autophagy for pestivirus infection. Autophagy was reported to be important for classical swine fever virus (CSFV) release (36, 37), BVDV and CSFV replication (38–42), as well as the regulation of the innate immune response in CSFV and BVDV infection (43–45). The importance of ATG14, ATG13, and ATG9A in the context of BVDV infection and autophagy was specifically reported (46, 47). A special role in the interference with autophagy was assigned to the NS4B protein of BVDV (48) and the NS5A protein of CSFV, which induces mitophagy (49). In this study, a selective disadvantage of mutations in the autophagy proteins ATG4B, ATG7, ATG3, and ATG12 could be identified, whilst mutagenesis of the VMP1 gene resulted in a strong selective advantage (Fig 2C). Therefore, our study provides further insights into the importance of individual genes involved in autophagy for BVDV infection.

VMP1 is a critical component of the autophagy initiation machinery across mammalian species (Sup Fig 2). In earlier CRISPR-Cas9 loss of function screens for resistance of human cells to infection with Zika and Yellow Fever Flaviviruses VMP1was identified as critical for survival (3). We also found VMP1 among the most selected genes in the population challenged with C87_mCherry-E2_, even though it is considered to be essential for viability. Re-testing of individual *VMP1* knockouts confirmed the key role of this protein for infection with both BVDV strains, as illustrated by powerful selection of *VMP1* mutants (Fig 4B). BVDV challenge led to an increase in the proportion of the two different *VMP1* mutants from practically undetectable to more than 20% under infection imposed selective pressure. Interestingly, infection with the NADL strain resulted in selection of a higher proportion of *VMP1* mutants suggesting existence of viral strain specific differences in requirement for VMP1 function.

We have noted that the same study that identified critical roles for VMP1 and TMEM41B (3) also determined that ATG7 was specifically not required for resistance to Flavivirus infections. This contrasts with our finding that *ATG7* mutants were preferentially depleted from BVDV challenged populations (Figs 2B and Table 2). One explanation for this discrepancy is that the previously published screen focused on positive selection of mutants while in our screen we attempted to capture both the positively and negatively selected knockouts by analyzing an early, day six timepoint. In fact, we found virtually no significant negative hits in our day eleven populations, presumably due to massive cell death at that time point that masks differences in the rate of depletion. It is also possible that the effect of ATG7 cannot be detected by comparison of pure mutant and control cell populations without prior identification of the precise MOI and timepoint that would produce measurable differences. In that respect, our approach of analyzing mixed population provides a continuous readout that can be precisely quantified.

For validation of hits generated by our screen we chose to analyze mixed populations of cells that are typically produced by transducing guide carrying lentiviruses at low MOI. We reasoned that such incomplete editing in fact provides an excellent platform for quantitative measurement of the effect of individual gene deletion on resistance to BVDV infection. Recently developed software tools allow deconvolution of sequencing traces of the edited locus to provide quantitative estimation of the number and types of mutations in the cell populations. We have successfully used these tools to analyze changes in the proportion of mutant cells following challenge with BVDV and confirmed that genes identified in the screen indeed change resistance of MDBK cells to BVDV infection. Furthermore, our approach has illustrated its suitability for analysis of the effect of deleterious mutations, like those that prevent creation and maintenance of sufficient numbers of cells to do comparative challenges.

Even though PHGDH phosphoglycerate dehydrogenase was not at the top of the list of the most selected mutants, we chose to include this gene for follow up tests. This decision was based on a recent report of this gene controlling CSFV infection (33). It has been shown that CSFV inhibits serine mediated antiviral immunity by inducing deacetylation of PHGDH, an enzyme that acts as a rate-limiting step in biosynthesis of serine. Because there were no reports linking PHGDH function to resistance to BVDV, we chose to test if the mutants in this gene indeed protect host cells against infection, as suggested by our screen results. Consistent with its role in the control of CSVF infection, we found that loss of PHGDH function promotes survival of BVDV infected cells as illustrated by a nearly 1.5-times increase in the proportion of *PHGDH* mutants in BVDV challenged populations of MDBK cells (Table 2). This observation suggests that PHGDH is a general factor controlling viral infections across mammalian species.

Overall, our study made several important steps towards developing novel methodology to study specifics of bovine host-pathogen interactions. It demonstrated the role of several genes that have previously not been linked directly to the control of BVDV infection and provided additional evidence supporting the existence of conserved viral resistance mechanisms.

## Materials and Methods

### Cell culture

HEK293T cell line used for packaging of lentiviruses was obtained from ATCC (CRL-1573), BVDV free MDBK cell (ATCC CCL-22) line maintained at the University of Veterinary Medicine, Vienna was used in all single gene knockout tests. Cells were maintained in DMEM (Merck) supplemented with 10% of BVDV-free Fetal Bovine Serum and 1x penicillin-streptomycin (ThermoFisher Scientific) at 37°C and 5%CO_2_.

### Library design

All possible 20 nucleotide long sequences adjacent to an NGG Cas9 PAM sequence were identified within all coding and a subset of non-coding transcripts annotated in the Ensembl bovine genome assembly ARS-UCD1.2 v104 (NCBI RefSeq GCF_002263795.1). Cutting efficiency for guides targeting coding genes was calculated using the complete Doench scoring model that considers guide sequence features as well as the location of the cut in the protein coding portion of the sequence (26). For each guide considered for inclusion in the library its off-target editing efficiency was estimated by calculating a “threat matrix” that was then used to calculate an individual off-target score. For each gene, individual guides were ranked based on the on-target cutting efficiency, off-target score and location of the cut within the coding sequence. The top 5 guides for each coding gene and lncRNA transcripts were included in the library. Following assembly, the library was further filtered to remove redundant sequences. The final library contained 114,024 unique elements targeting 22,964 genomic features of which 21,484 were annotated protein coding genes as well as 1,000 control non-targeting guides. The library was synthesized by GENEWIZ and cloned into BbsI sites of a custom PLV2.8 vector that was created in house by replacing the puromycin resistance gene in pKLV2-U6gRNA5(BbsI)-PGKpuro2ABFP-W (Addgene 67974) with the sequence of the *BSD* gene coding for blasticidin resistance. Representation of guides in the final cloned library was confirmed by Illumina sequencing.

### Inducible CRISPR-Cas9 MDBK cell line

5×10^6^ MDBK TetOn cells were transfected by electroporation employing the Amaxa device, program X-001, with 0.25 µg of plasmid pEF-PAC (encoding a puromycin resistance, linearized with HindIII) and 2.5 µg of an *in house* generated plasmid derived from pTRE expressing Cas9 with a C-terminal FLAG tag followed by a P2A cleavage site and GFP under the control of the tetracycline responsive element (linearized with PvuI). The cells were plated on a 10cm dish, and 48h after transfection the medium was changed to include 1µg/ml puromycin to select for positive clones. Once single colonies appeared, they were picked and each transferred into an individual well of a 96 well plate. The resulting clones were validated by testing for expression of GFP following exposure to 2.5 µg/ml doxycycline. The efficiency of knock-out mediated by sgRNAs targeting either GFP or the host essential gene myc was subsequently tested. MDBK_CAS9_ cells were transduced with lentiviral vectors encoding a sgRNA targeting GFP. Subsequently, GFP specific fluorescence was monitored for 7 passages by flow cytometry in comparison with MDBK_CAS9_ cells transduced with a control plasmid. To also test the KO efficiency for an endogenous gene, MDBK_CAS9_ cells were transduced with lentiviral vectors encoding a sgRNA targeting *myc* and mCherry. Subsequently, the number of mCherry expressing cells was monitored for seven passages by flow cytometry and compared with the number of mCherry expressing cells in the control condition transduced with an mCherry expressing control plasmid. The susceptibility of MDBK_CAS9_ cells to infection with BVDV was also evaluated in comparison to parental MDBK cells using an mCherry labelled BVDV strain C87_mCherry-E2_ clone.

### Lentiviral transduction

Lentiviral library was introduced into MDBK cells by spinfection following using optimized Broad institute protocol (https://portals.broadinstitute.org/gpp/public/resources/protocols). Briefly 1×10^8^ cells were centrifuged at 920g for 2 hours at 30°C in 6 well plates and placed into 37°C 5%CO_2_ incubator overnight. Next day all cells were transferred to T175 flasks at 1×10^7^ seeding density. After 48 hours 2.5 µg/ml of doxycycline and 10 µg/ml of blasticidin were added to each flask. 24 hours later doxycycline containing media was removed, and after a wash with DPBS replaced with fresh media supplemented to 10 µg/ml blasticidin. Cells were grown with continuous selection for 8 additional days and split twice prior to shipment to Vienna.

### BVDV challenge and gDNA extraction

CRISPR knockout cell lines were seeded upon arrival in Vienna and after one passage they were trypsinized and counted with a DeNovix cell counter. 1×10^6^ cells each were seeded on 6-well plates in Dulbecco’s Modified Eagle Medium (DMEM, L0104-500, Biowest) with 2 µg/mL puromycin, 1x Penstrep and 10% BVDV-free batch of fetal calf serum (Corning, 35-079-CV). Cells were challenged using a MOI of 0.1 of two different cytopathogenic BVDV-1 virus strains, C87_mCherry-E2_ and NADL. Unchallenged cells served as negative controls. As soon as confluency reached <50% on the tissue culture plate, typically 5-9 days post infection, cells were trypsinized and genomic DNA was extracted following the manufacturer’s protocol with a Monarch gDNA kit (NEB, T1020L). gDNA of cells staying above the 50% confluency threshold was isolated on day 9-14. Negative control cells were taken +/-1 day in parallel for extraction. The gDNA amount of the samples was quantified with a QuantiFluor® ONE dsDNA System (Promega, E4871) prior to shipping and sequencing steps.

### Guide abundance quantification

Amplicon libraries for next generation sequencing were generated by PCR amplification of genomic DNA templates using Broad Institute hybrid primers composed of Illumina cell attachment and sequencing primer sites as well as lentiviral vector sites flanking the guide sequence P5 Argon and P7 Beaker. Absolute abundance of individual guides in the raw sequencing data of amplicons recovered from the library transduced cells that were unchallenged or exposed to various strains of BVDV was quantified using the *count* function of the MAGeCK Bioconductor package (43). Cross-sample abundance values were normalized using the 1000 non-targeting control gRNAs.

### Generation of single gene knockout MDBK cell lines

For a subset of genes that were positively and negatively selected in the screen, two CRISPR Cas9 targeting guides (Table 1) were cloned into the lentiCRISPRv2-Opti vector (163126, Addgene). The targeting clones were validated by sequencing. Lentiviral particles were then generated according to Addgene protocol (http://www.addgene.org/tools/protocols/plko/#E). In brief, 2.5×10^6^ HEK 293T cells were seeded in 6-well plate. Cells were transfected the day after seeding with 1 μg of the psPAX2 plasmid (12260, Addgene), 0.67 μg of the pMD2.G plasmid (12259, Addgene) and 1.3 μg of the lentiCRISPRv2 sgRNA plasmids using GeneJuice transfection reagent (70967, Merck) according to manufacturer protocol. Media was refreshed 12–15 hours post-transfection. Virus-containing supernatant was collected 48hrs and stored at −20°C for further use. 1×10^6^ MDBK cells in 1.5 ml media were transduced with 0.5 ml virus-containing supernatant in the presence of 2 μg/ml of polybrene. Cells were incubated overnight and resuspended in fresh media containing 2 μg/ml puromycin. After 7 days of selection, cells were challenged with BVDV strains NADL and C87_mCherry-E2_ for 6 to 11 days.

### Gene editing analysis

Genomic regions flanking the predicted cut site for each single gene targeting experiment were amplified using a pair of primers listed in Sup Table 2. PCR products were sequenced using an internal primer listed in Sup Table 2. Resulting sequencing traces were deconvoluted using locally run *ice* software (34).

---

**Sup Fig 1.**
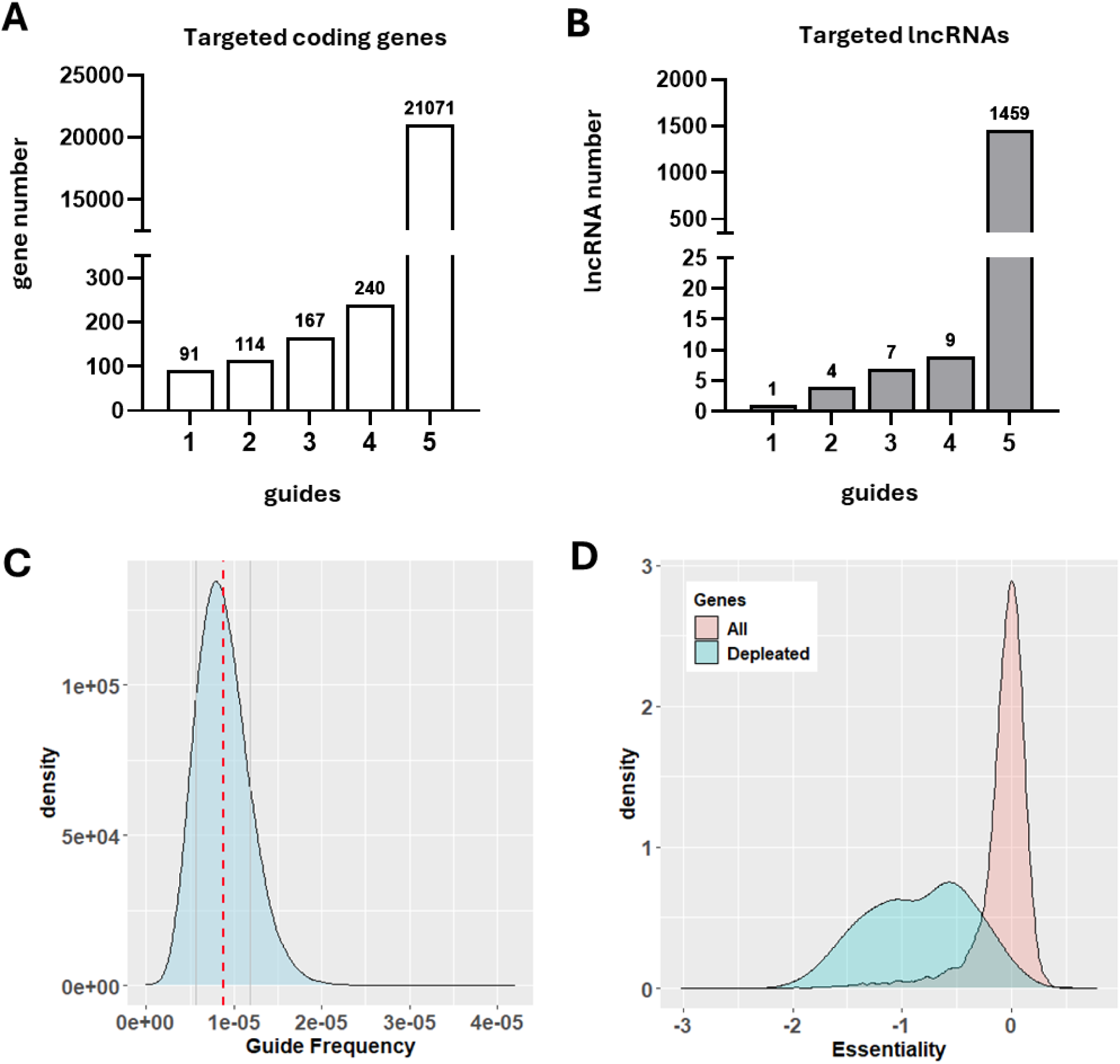
Bovine whole genome knockout library metrics. The majority of annotated bovine coding genes (**A**) and lincRNAs(**B**) have 5 guides each included in the library. The final cloned library shows uniform representation of individual guides centered around the value of expected frequency (dashed red line) with a majority (70% of guide frequencies found within 1 SD from the expected value (**C**). Distribution of compound human kidney cells essentiality scores for all mapped bovine homologues (red) and 352 mutants that were significantly depleted by Day 22 of MDBK cell culturing (blue) (**D**).

**Sup Fig 2.**
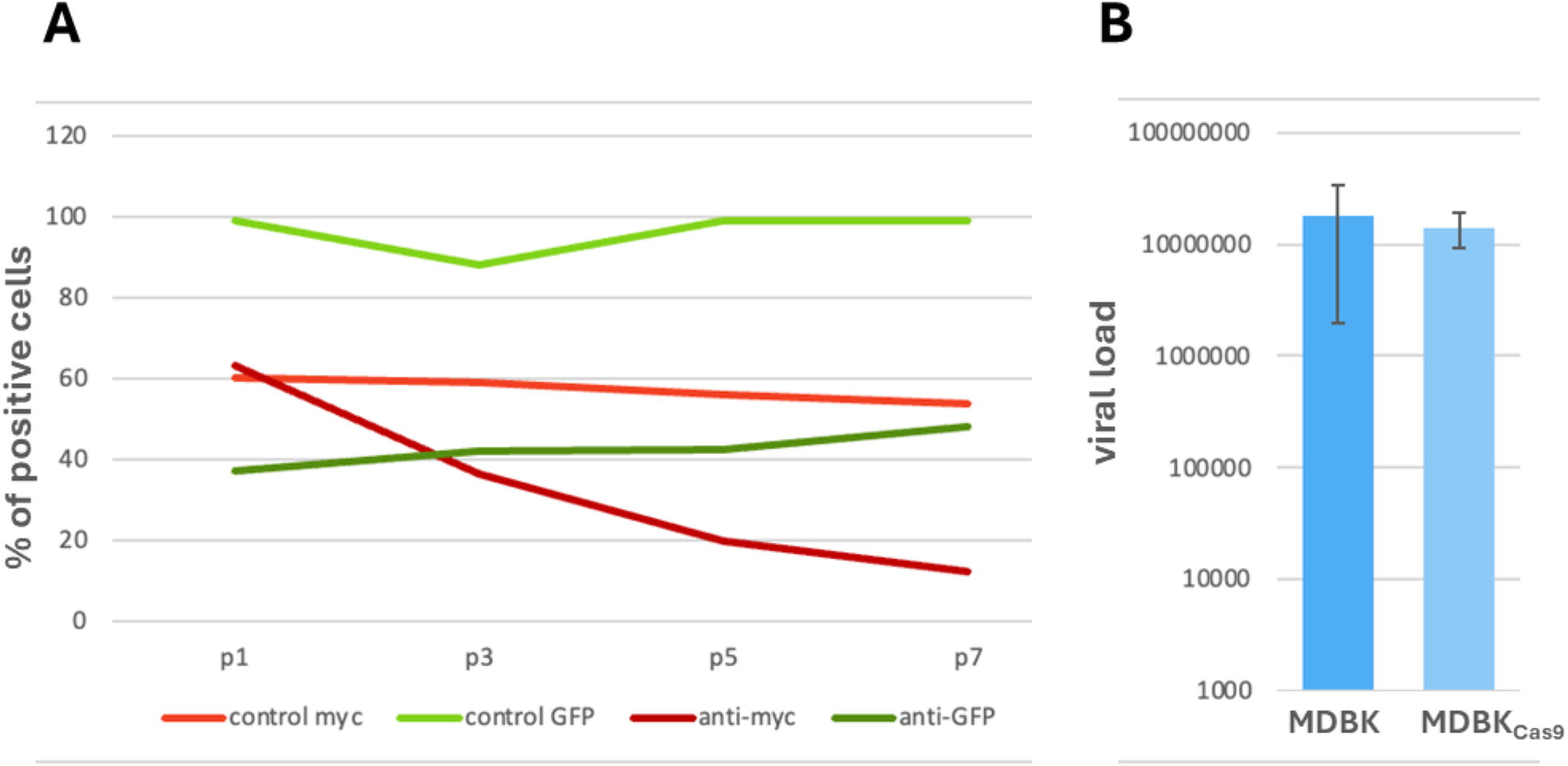
Characterization of Cas9 expressing MDBK cell line. **A.** MDBK_Cas9_ cells transduced with guides targeting GFP (dark green) or *myc* (dark red) show dramatic decrease in the levels of respective fluorescent proteins, as measured by flow cytometry for 7 passages (24 days). **B.** MDBK_Cas9_ and parental MDBK cell lines are infectible by BVDV.

**Sup Fig 3.**
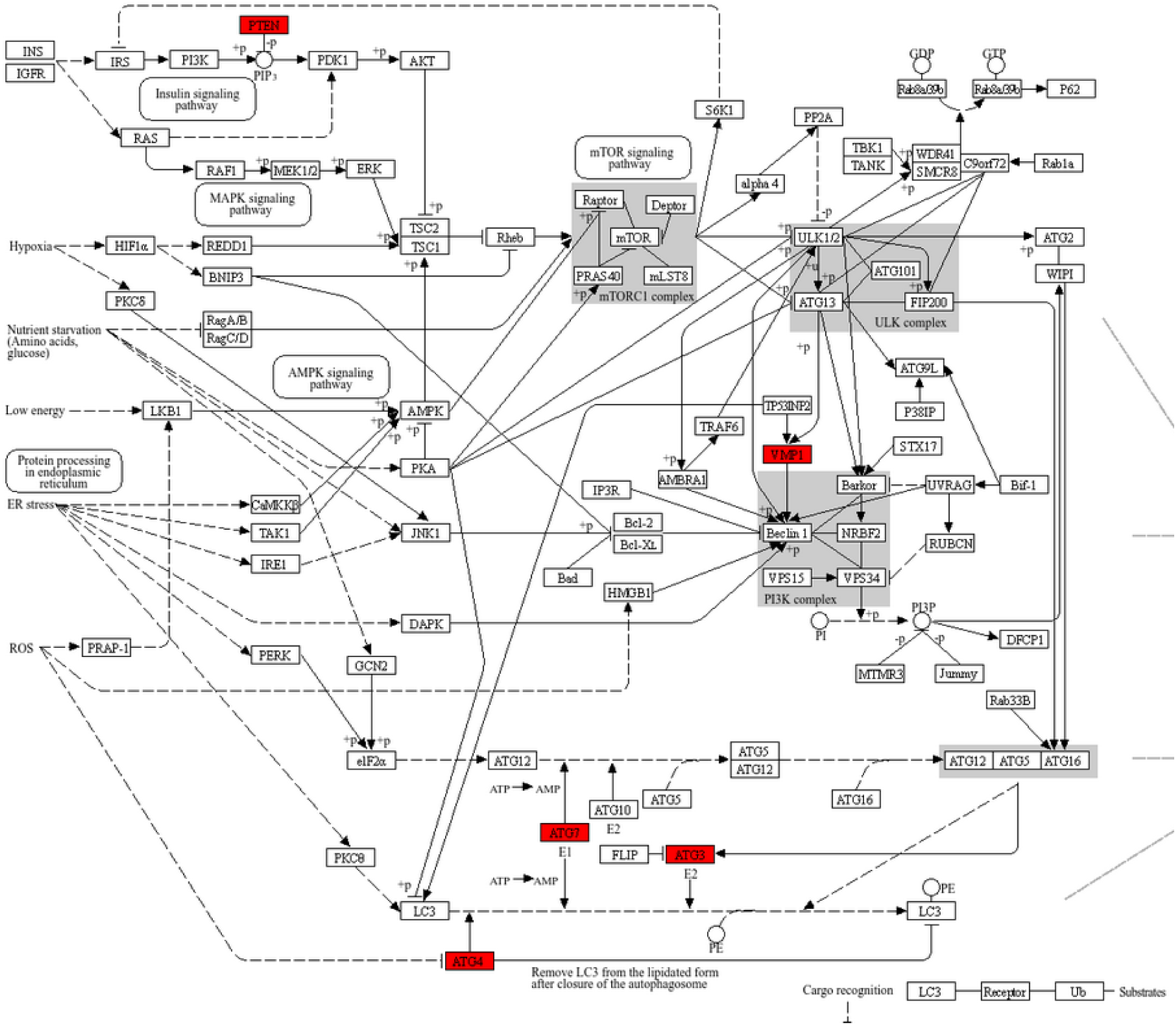
Animal autophagy KEGG pathway showing genes identified as differentially selected after six days following C87_mCherry-E2_ challenge of MDBK cells

**Supplementary Table 1.**
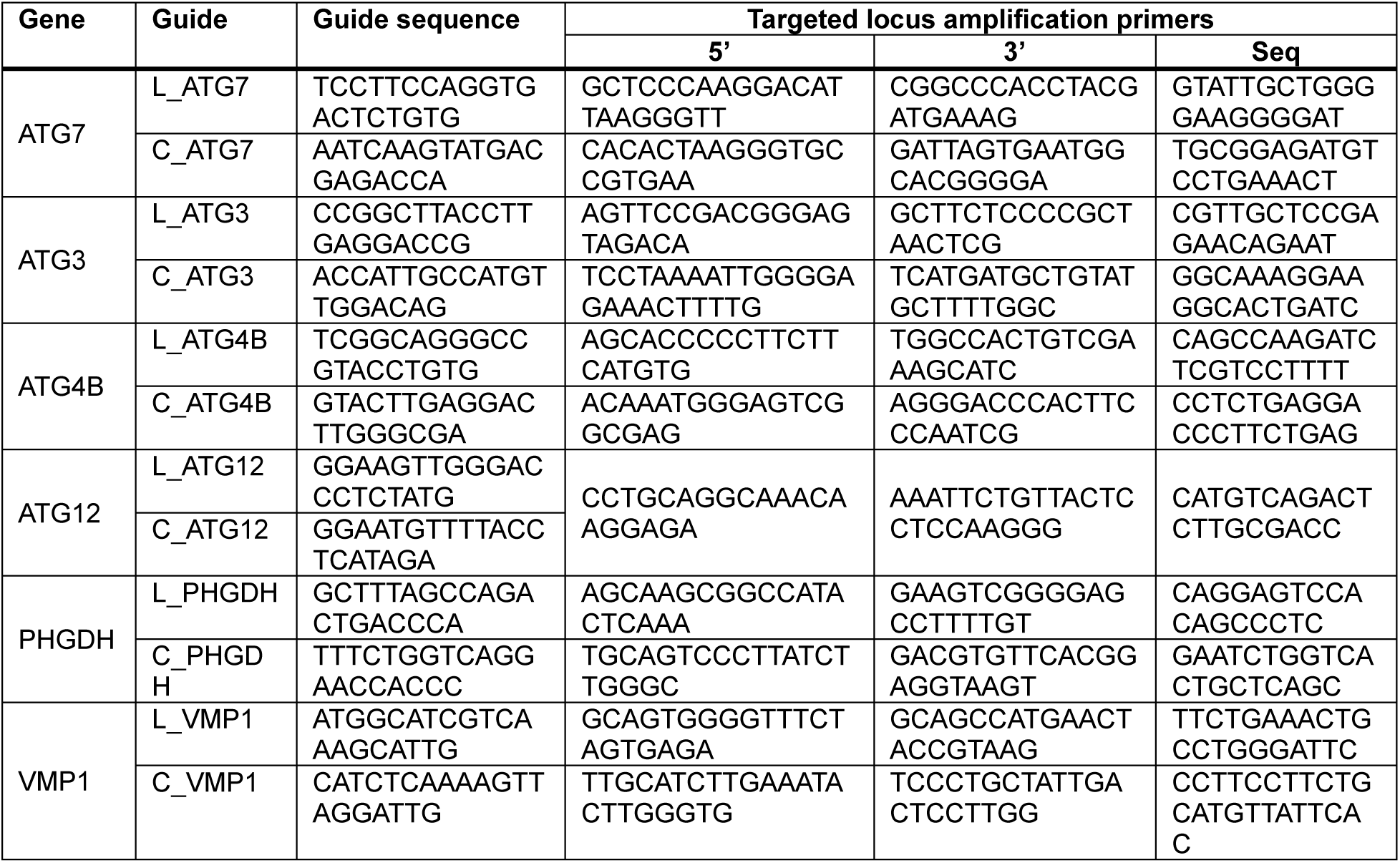
Sequences of sgRNA guides and locus specific primers used in deconvolution analysis of single gene knockouts.

**Supplementary Table 2.** Gene knock outs highly enriched (FDR<0.05) by Day 11 of BVDV infection.

| **pCR464** | **Common** | **NADL** |
| --- | --- | --- |
| CRLS1 | ADAM17 | VPS16 |
| AGO2 | RHBDF2 | TMEM41B |
| PKMYT1 | RABGEF1 | MOGS |
| THEM6 | SRRD | PPP1R21 |
| SYVN1 | HSPA13 | PTPN23 |
| TSC1 | RBSN | COL10A1 |
| SEC11A | SPCS3 | PPP1R8 |
| VMP1 | VPS45 | VPS8 |
| YIPF5 |  | VPS11 |
| SPRYD7 |  | RPS3 |
| NOB1 |  | SCAMP2 |
|  |  | POLR1A |
|  |  | KPNB1 |
|  |  | PRKCSH |
|  |  | HM13 |
|  |  | TMCO1 |
|  |  | ARHGEF40 |
|  |  | CAPRIN1 |
|  |  | GANAB |
|  |  | PROCA1 |
|  |  | DROSHA |
|  |  | PTPMT1 |
|  |  | TSC2 |
|  |  | ANKFY1 |
|  |  | RAB5IF |
|  |  | ELL3 |
|  |  | UBE2G2 |

## Acknowledgments

This work was supported by Norwegian Research Council grants 281928 and 309707 and Austrian Science Fund (FWF) P35674

